# Targeted degradation of endogenous condensation-prone proteins improves crop performance

**DOI:** 10.1101/2024.11.13.623495

**Authors:** Ming Luo, Qing Wen, Sitao Zhu, Hua Dang, Ruixia Niu, Jiawei Long, Zhao Wang, Yongjia Tong, Yuese Ning, Meng Yuan, Guoyong Xu

## Abstract

Effective methods, such as CRISPR and RNA interference, exist for modulating gene expression at DNA and RNA levels, but approaches for directly modifying endogenous proteins remain lacking in plants. Here, we develop a targeted condensation-prone-protein degradation (TCD) strategy to eliminate endogenous proteins, particularly those prone to condensation. We identify an E3 ligase, E3TCD1, that degrades itself but selectively targets other proteins when fused to them. In rice, transgenic E3TCD1 fusions with Teosinte branched 1 and Early flowering 3 (OsELF3) modulate tiller numbers and flowering times, respectively. The TCD system is also controllable. Using the Pro_TBF1_-uORFs_TBF1_ expression control cassette, we can conditionally degrade the negative defense regulator OsELF3 upon pathogen invasion, enhancing rice resistance without interfering with rice flowering time. This method, unlike animal-targeting strategies, does not rely on small molecules, antibodies, or genetic knock-ins, showing promise as a gene therapeutic avenue for optimizing crop performance and potentially addressing human diseases.

## Introduction

Plants dynamically regulate their proteostasis, encompassing protein synthesis, modification, trafficking, localization, and degradation, to adapt to diverse growth requirements and navigate biotic and abiotic stresses. Recent investigations highlight the significance of protein condensation through phase separation and transition as an additional critical layer in proteostasis regulation ^1–7^. This layer plays a substantial role in shaping plant development and responding to stressors ^8–30^. Thus, modulating proteostasis is a promising breeding strategy that could complement well-established genetic engineering techniques like CRISPR-Cas and RNA interference, which operate at the DNA or RNA levels to enhance plant performance ^31–33^. However, methods for applying such a strategy in plants remain largely unexplored.

In the intricate proteostasis regulatory network, the ubiquitin-proteasome system (UPS) and the autophagy system are recognized as integral components ^34–36^. Tools based on these two systems, like the auxin-inducible degron, autophagy-targeting chimeras (AUTACs), PROteolysis TArgeting Chimeras (PROTACs), and Trime have proven invaluable for human basic research and drug discovery by effectively targeting protein for degradation ^37–40^. However, the application of these systems in agriculture faces challenges, as they often depend on small chemical molecules, antibodies, or the genetic knock-in of degron sequences fused to endogenous proteins, rendering them time-consuming and expensive ^41^. Hence, developing a straightforward and effective method for directly modifying endogenous plant proteins is essential for fundamental research and practical applications in agriculture. Such an approach holds promise for rapid genetically improved crop performance, aligning with the broader goals of sustainable agriculture.

## Results

### Screen E3 for the targeted condensation-prone-protein degradation system (TCD)

A hallmark of protein condensation is the remarkable phenomenon of large-scale coalescence, where identical molecules come together from the soluble phase to form smaller-sized complex and larger-sized condensates ^4,42,43^. We propose a genetically engineered protein to mimic the endogenous target protein (X), sneak into X-containing condensates and trigger protein degradation (Fig. 1a). We assumed that fusing the X protein with an E3 ligase (X–E3) from the UPS system would bring it in proximity to X-containing condensates through the X protein. We opted for Really Interesting New Gene (RING) E3 ubiquitin ligases due to their diversity and ability to function in a monomeric form in plants ^44,45^. To promote the incorporation of more identical X–E3 proteins into the targeted condensates, we selected E3 ligase to be intrinsically disordered proteins (IDPs) prone to cluster ^4,42,43^. We predicted 51 out of 508 possible RING E3 in Arabidopsis as IDPs (E3IDP1–51) and successfully cloned 44 for further screening (Extended Data Fig. 1a; Supplementary Table 1). As protein concentration increases, it promotes protein condensation ^43^. Therefore, we overexpressed these proteins tagged with the yellow fluorescence protein (E3IDP–YFP) in *N. benthamiana* to determine their condensation behaviors. We discovered that 17 RING E3 proteins formed visible condensates (e.g., E3IDP1), while 17 did not (e.g., E3IDP6; Extended Data Fig. 1a, b). Surprisingly, ten E3 ligases did not show obvious YFP signals, but the co-expressed CFP control was always detected in the nucleus (e.g., E3IDP45; Extended Data Fig. 1a, c). We hypothesized that some E3 ligases could recognize themselves as substrates and constitutively degrade themselves.

**Fig. 1.**
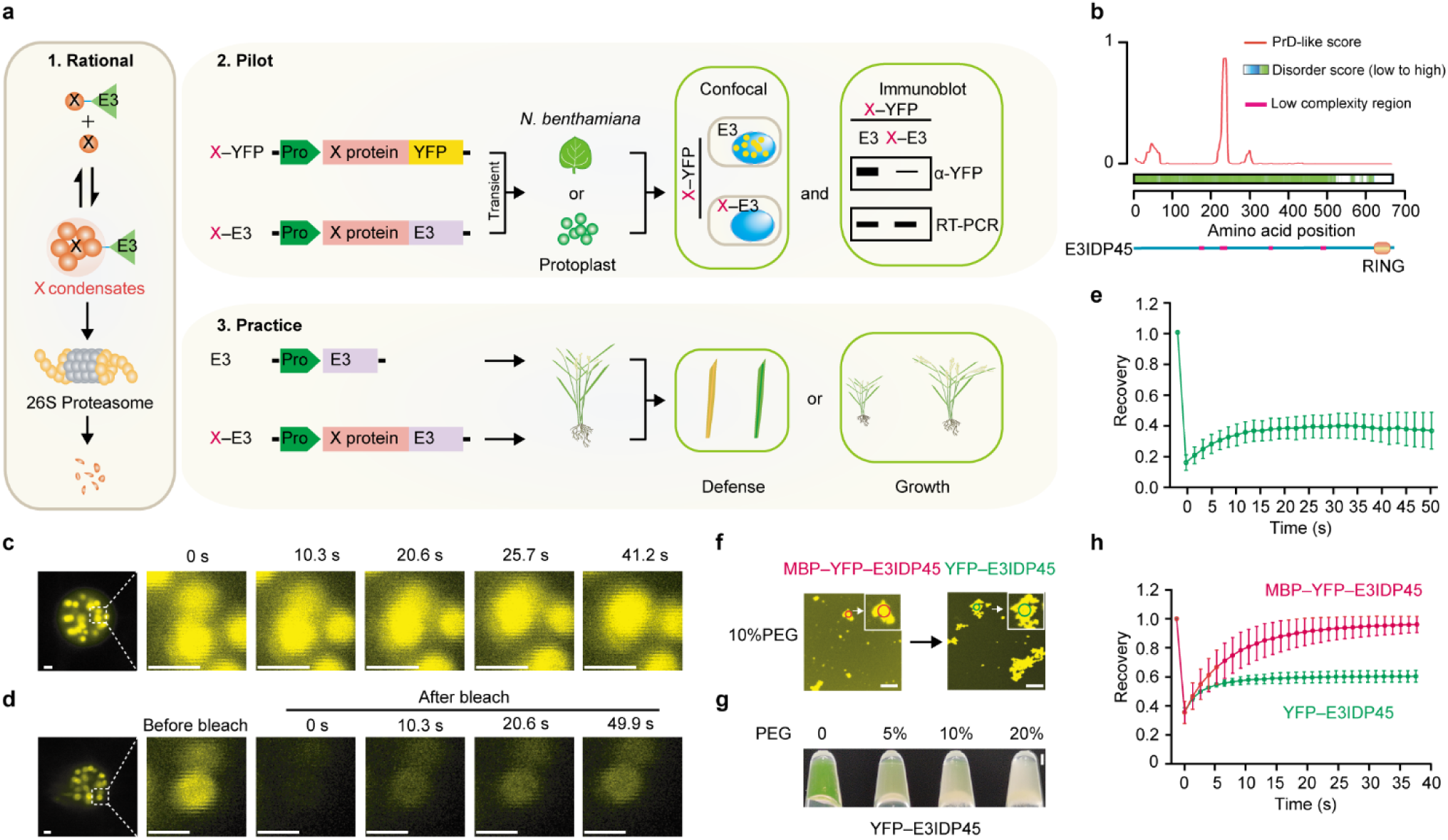
Screening E3IDP45 (E3TCD1) for the targeted condensate degradation (TCD) system. See also Extended Data Fig. 1. **a**, The schematic diagram of TCD. The TCD system targets specific condensates formed by the condensation-prone protein X. These proteins typically aggregate into condensates, transitioning from soluble phases to organized structures, achieving equilibrium across different scales of condensates (oligomers, smaller-sized and larger-sized). To remove such condensates, an E3 ligase from the ubiquitin-proteasome system (UPS) is fused as X–E3. This fusion protein mimics X, aiding its penetration into the condensate. Once inside, X–E3 triggers the degradation. Before generating transgenic plants for phenotyping, pilot experiments involving transient expression of X–E3 with X–YFP in *N. benthamiana* or protoplasts are crucial to assess TCD efficiency via methods including microscopic observation or immunoblot analysis. Different promoters are necessary to control the spatial and temporal expression as well as the expression intensity of X–E3. **b**, Prediction of E3IDP45 as an intrinsically disordered protein (IDP) using PLAAC and D2P2 algorithms indicated by Prion domain (PrD)-like score and disorder score, respectively. The bottom section displays low complexity regions and the RING domain as annotated by the website SMART. **c**, Fusion behavior of E3IDP45^ΔRING^–YFP condensates after transient expression in *N. benthamiana*. E3IDP45^ΔRING^, E3IDP45 without the RING domain. **d**, **e**, Fluorescence recovery after photobleaching (FRAP) of E3IDP45^ΔRING^ condensates. Scale bar, 2 µm. **f**, In vitro condensation of YFP–E3IDP45 in the presence of the crowding agent PEG before and after cleavaging the molecular chaperon, maltose-binding protein (MBP). Scale bar, 20 µm. **g**, YFP–E3IDP45 condensation in different concentrations of PEG. Scale bar, 1 mm. **h**, FRAP of the fibrous condensates circled in (f). The points and error bars show the mean ± s.d. of the time course of the recovery after photobleaching in (e, h, *n* = 5).

To validate our hypothesis, we experimented with these ten E3 by deleting the predicted RING domain (E3IDP^ΔRING^; Supplementary Table 1), which plays a crucial role in recruiting other components of the UPS system. We observed that five E3 ligases failed to generate YFP signals after removing the predicted RING domains (e.g., E3IDP8; Extended Data Fig. 1c). However, we observed the reappearance of YFP signals as condensates for the remaining five E3 after removing the predicted RING domains (e.g, E3IDP45; Extended Data Fig. 1c). Thus, these E3s are the best candidates for engineering X–E3 fusions because they may be self-degraded and not remain in cells before and after clearing X-containing condensates.

### Identifying E3IDP45 for the TCD system (E3TCD1)

Subsequently, we focused on one of the five candidates, E3IDP45, which shares a typically low-complexity region with the other candidates (Fig. 1b and Extended Data Fig. 1d). Upon transient expression in *N. benthamiana*, we observed the fusion behavior among the E3IDP45^ΔRING^ condensates (Fig. 1c). Furthermore, fluorescence recovery after photobleaching (FRAP) revealed consistent interchangeability of these condensates with the surrounding soluble fractions, indicative of liquid-liquid phase separation (Fig. 1d, e). As we focused on utilizing these E3IDPs for engineering purposes, we did not assess their potential for condensation at the natural expression level using the native promoters. Instead, we conducted further analysis of its condensation behavior in vitro.

We expressed maltose-binding protein (MBP)-tagged YFP–E3IDP45 in *Escherichia coli*. Initially, before removing MBP, we observed that the purified MBP–YFP–E3IDP45 exhibited predominantly spherical droplets with fewer fibrous condensates when exposed to a crowding agent that promotes intermolecular interaction (Fig. 1f). However, upon removal of the molecular chaperone MBP, YFP– E3IDP45 underwent a rapid phase transition, resulting in increased turbidity with escalating concentrations of the crowding agent (Fig. 1g). Notably, we observed swift formation of non-liquid fibrous condensates emerging from the spherical droplets (Fig. 1f). These fibrous condensates displayed reduced material exchange with the surrounding solution, as evidenced by their slower recovery rate after bleaching compared to those formed by MBP–YFP–E3IDP45 (Fig. 1h). Thus, our findings indicate that E3IDP45 exhibits a high propensity for condensation. These findings indicate that the E3IDP45 can recognize itself as a substrate for degradation, rendering it well-suited for engineering the TCD system. Such E3 ligases that meet the requirements of the TCD system are named with the suffix E3TCD. Therefore, E3IDP45 is renamed as E3TCD1.

### Cross-activity examination of the TCD system

To validate the TCD system, we first implemented the TCD experimental setup using E3TCD1^ΔRING^ as the X protein to examine whether E3TCD1 could eliminate E3TCD1^ΔRING^. Once again, we failed to detect E3TCD1–YFP through microscopic observation or immunoblot analysis (Fig. 2a, b). Subsequently, we expressed E3TCD1^ΔRING^–YFP and observed the re-emergence of condensates in the nucleus (Fig. 2a). Remarkably, co-expression of the full-length E3TCD1 as X–E3 effectively diminished these E3TCD1^ΔRING^–YFP condensates (Fig. 2a). Immunoblot analysis confirmed the depletion of E3TCD1^ΔRING^–YFP at the protein level, while their mRNA levels remained consistent regardless of co-expression with E3IDP45 (Fig. 2b).

**Fig. 2.**
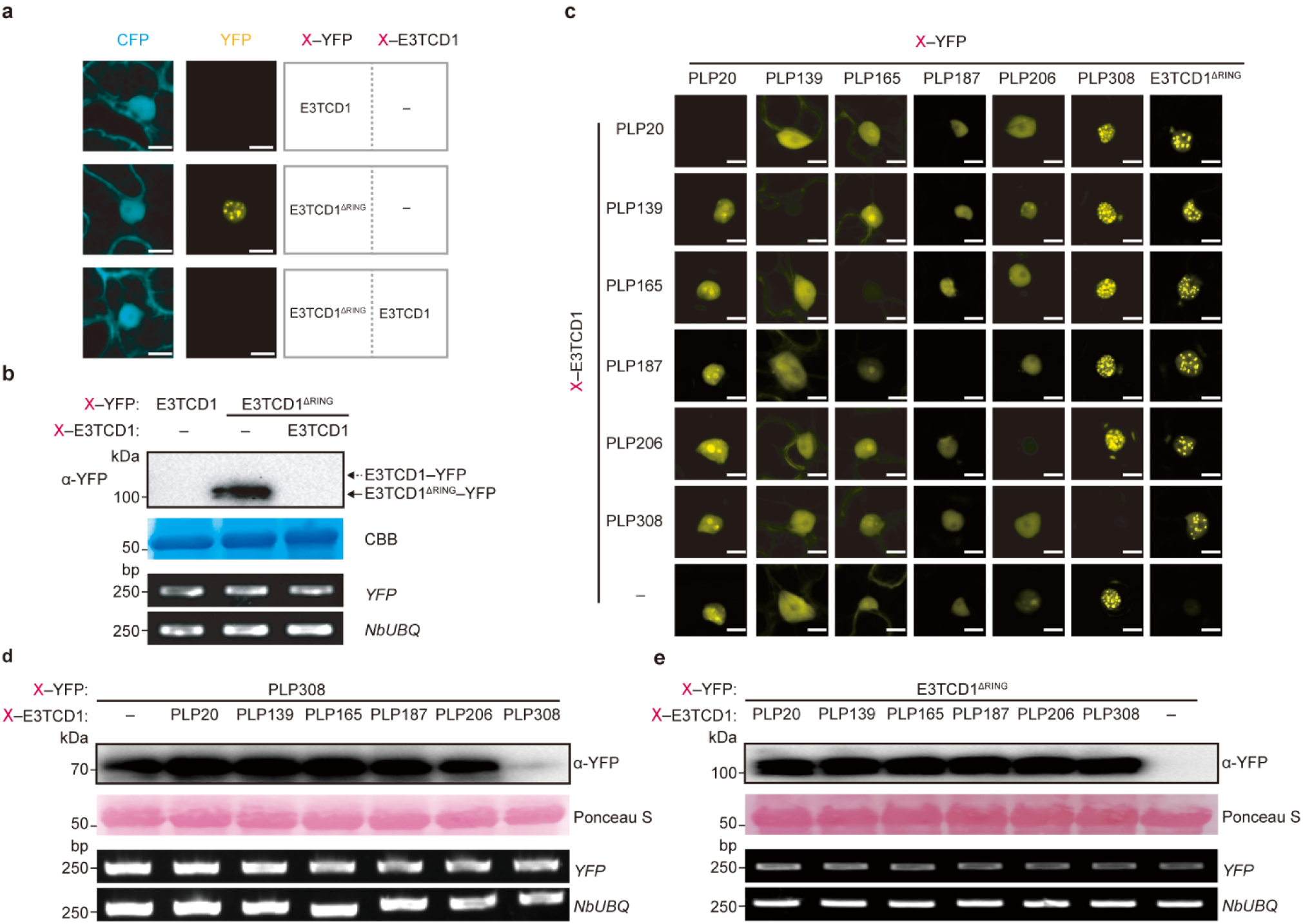
Cross-activity examination of the TCD system. See also Extended Data Fig. 2. **a**, **b**, Degradation of E3TCD1– YFP and E3TCD1^ΔRING^–YFP condensates by E3TCD1 in the pilot experiments of microscopic observation (a) and immunoblot analysis (b) in *N.benthamiana*. The target protein X is tagged with YFP, while the TCD is indicated by X– E3TCD1. For microscopic observation, 35S::CFP serves as the control (a). Scale bar, 10 µm. CBB, Coomassie brilliant blue. **c**, Microscopic observation of each X–protein after co-expression with different E3TCD1 fusion proteins (X– E3TCD1). Six transcription factors and the E3TCD1^ΔRING^ were examined as the target X protein. Scale bar, 10 µm. **d**, **e**, Immunoblot analysis of PLP308–YFP (d) and E3TCD1^ΔRING^–YFP (e) after co-expression with different E3TCD1 fusion proteins (X–E3TCD1). Semi-quantitative RT-PCR is conducted against YFP with *NbUBQ* as the internal control (b, d, e).

To further validate the TCD system, we focused on two key questions: First, can fusing E3TCD1 with various X proteins effectively induce their degradation? Second, is this degradation specific to the X proteins? For the first question, we specifically chose transcription factors (TFs) for examination due to mounting evidence across animal and plant studies indicating that TFs and their regulators form biomolecular condensates to regulate transcription ^8,10,25–27,46–50^. We fused E3TCD1 to eight TFs from the TCP, AP2, bZIP, ZnF and NAC families of rice (Supplementary Table 2) to generate TF–E3TCD1 constructs, which were co-expressed with their corresponding TF–YFP in *N. benthamiana*. Our findings revealed that six of the TF–E3TCD1 fusion proteins were capable of degrading their respective TF–YFPs, as indicated by reduced fluorescence (PLP20/139/165/187/206/308; Extended Data Fig. 2a). We successfully conducted immunoblot analysis on six out of the eight TF–YFP samples and found results consistent with our fluorescence observations (five shown in Extended Data Fig. 2b with PLP308 shown below).

In addition, we noted that the TCD mechanism was effective not only in degrading proteins capable of forming condensates (PLP20/165/206/308) but also in those existing in the soluble fraction (PLP139/187). Given that all TFs are predicted to possess disordered regions (Extended Data Fig. 2c), it is plausible that the diffused TFs may cluster in sizes below the detection limit or necessitate additional stimulating conditions for visible condensate formation, as previously suggested ^13,16,17^. Notably, two TFs were not efficiently degraded in these assays (PLP32/247; Extended Data Fig. 2a). While the exact system requirements for effectiveness were not fully established, we strongly recommend rapid pilot experiments involving microscopic observation and immunoblot analysis before employing TCD for degrading the target X protein in subsequent studies (Fig. 1a).

For the second question, we tested each TF–E3TCD1 fusion with the six degradable TFs and revealed selective degradation of the corresponding target only (Fig. 2c). Immunoblot analysis further confirmed that PLP308–YFP was exclusively degraded by PLP308–E3TCD1, not by other TF– E3TCD1 fusions or E3TCD1 alone (Fig. 2d). We also investigated whether the X–E3TCD1 fusion could recognize substrates of E3TCD1. Our observations, using both microscopy and immunoblot analysis, revealed that only E3TCD1 itself—but not the other six TF–E3TCD1 fusions—was capable of degrading E3TCD1^ΔRING^–YFP (Fig. 2c, e). While these results are robust and suggest that X– E3TCD1 fusions likely do not recognize substrates of E3TCD1, we have not fully explored all possible unidentified substrates of E3TCD1. We thus suggest using E3TCD1 as a control in pilot or subsequent comparison experiments, though it may not be strictly necessary. These findings highlight the efficiency and specificity of the TCD system in degrading endogenous proteins.

### Genetic analysis with the TCD system

We aimed to investigate whether the degradation of endogenous proteins via the TCD system induced any phenotypic changes. To assess the TCD system’s impact, we employed both transient and transgenic methods, which are commonly utilized for determining protein functions. Our transient experiment focused on PLP308, a rice Zinc-finger TF whose physiological roles remain unknown. Co-expression of PLP308–YFP with the control E3TCD1 in *N. benthamiana* led to noticeable macroscopic cell death, as evidenced by intensified trypan blue staining and elevated ion leakage (Extended Data Fig. 3a, b). Pilot experiments indicated reduced formation of PLP308–YFP condensates and protein levels upon co-expression with PLP308–E3TCD1 (Extended Data Fig. 3c, d).

As the co-expression assay primarily reflects the final outcome, the observed degradation efficacy might be attributed to the clearance of PLP308–YFP before they reached high expression levels. Consequently, we assessed whether PLP308–E3TCD1 could continue removing PLP308–YFP even after it accumulated to high expression levels and formed extensive condensates. We expressed PLP308–YFP for 48 hours and then induced PLP308–E3TCD1 expression using the β-estradiol-inducible system (Extended Data Fig. 3e). Remarkably, PLP308–E3TCD1 effectively degraded PLP308–YFP condensates even at high expression levels (Extended Data Fig. 3f, g). Furthermore, we found that the UPS system inhibitor, MG132, could inhibit PLP308–E3TCD1-triggered degradation of PLP308–YFP (Extended Data Fig. 3g). Therefore, co-expression with PLP308–E3TCD1 resulted in less pronounced cell death (Extended Data Fig. 3a, b). This suggests that PLP308–E3TCD1 initiates the degradation of PLP308–YFP through the TCD, thereby mitigating the associated phenotype.

To evaluate the effect of the TCD system in transgenic plants, our primary focus was on a rice TCP-type TF, Teosinte Branched 1 (OsTB1). This choice was motivated by the clear phenotype observed in rice with mutations in *OsTB1* DNA, which results in increased tillering numbers ^51–54^. OsTB1 has a predicted IDR (Extended Data Fig. 4a). We discovered an irregular condensate of OsTB1–YFP in the nucleus, indicating that OsTB1 has the potential to form condensates (Extended Data Fig. 4b). The pilot experiments showed a decrease in OsTB1–YFP condensates and protein after co-expressing OsTB1–E3TCD1, suggesting that OsTB1–E3TCD1 can degrade OsTB1 (Extended Data Fig. 4b, c). We introduced OsTB1–E3TCD1 (35S::OsTB1–E3TCD1) and the E3TCD1 control (35S::E3TCD1), both driven by the CaMV 35S promoter, into the rice cultivar Zhonghua 11 (ZH11). We found no statistically significant difference in the tillering number between ZH11 and the 35S::E3TCD1 control lines. This suggests that the self-degradation property of E3TCD1 prevents potential negative impacts on plant growth (Extended Data Fig. 4d, e). The transgenic lines with 35S::OsTB1–E3TCD1 showed increased tillering numbers compared to 35S::E3TCD1, consistent with plants that have the DNA mutation on *OsTB1*, as reported (Extended Data Fig. 4 d, e). We then performed RT-PCR against the untranslated region, which was not included in the OsTB1–E3TCD1 transgene, and detected similar mRNA levels for the endogenous *OsTB1* gene. This suggests that the phenotype was not caused by potential RNA interference (Extended Data Fig. 4f). These results suggest that the TCD system can efficiently degrade endogenous proteins and produce desirable traits.

### Using the TCD to modulate flowering time

The successful application of TCD in regulating rice tillering underscores the promising potential of this system for optimizing crop performance. Next, we focused on modulating flowering time, known as heading date, a crucial agronomic trait in rice that influences its region adaptation and grain yield. We targeted rice early flowering 3 (ELF3) because its *Arabidopsis* homolog (AtELF3) is a condensation-prone protein involved in transcriptional regulation ^10,55^. The pilot experiment in *N. benthamiana* showed that AtELF3–E3TCD1 reduced AtELF3–YFP condensates and protein, indicating its potential to degrade AtELF3 (Extended Data Fig. 5a, b). Again, the control line 35S::E3TCD1 did not negatively impact the plant growth compared to the non-transformed Col-0. However, transgenic plants expressing 35S::AtELF3–E3TCD1 displayed a notable tendency toward early flowering (Fig. 3a, b). This observation aligns with findings from plants harboring mutations in *AtELF3* DNA, as previously reported ^76,77^.

**Fig. 3.**
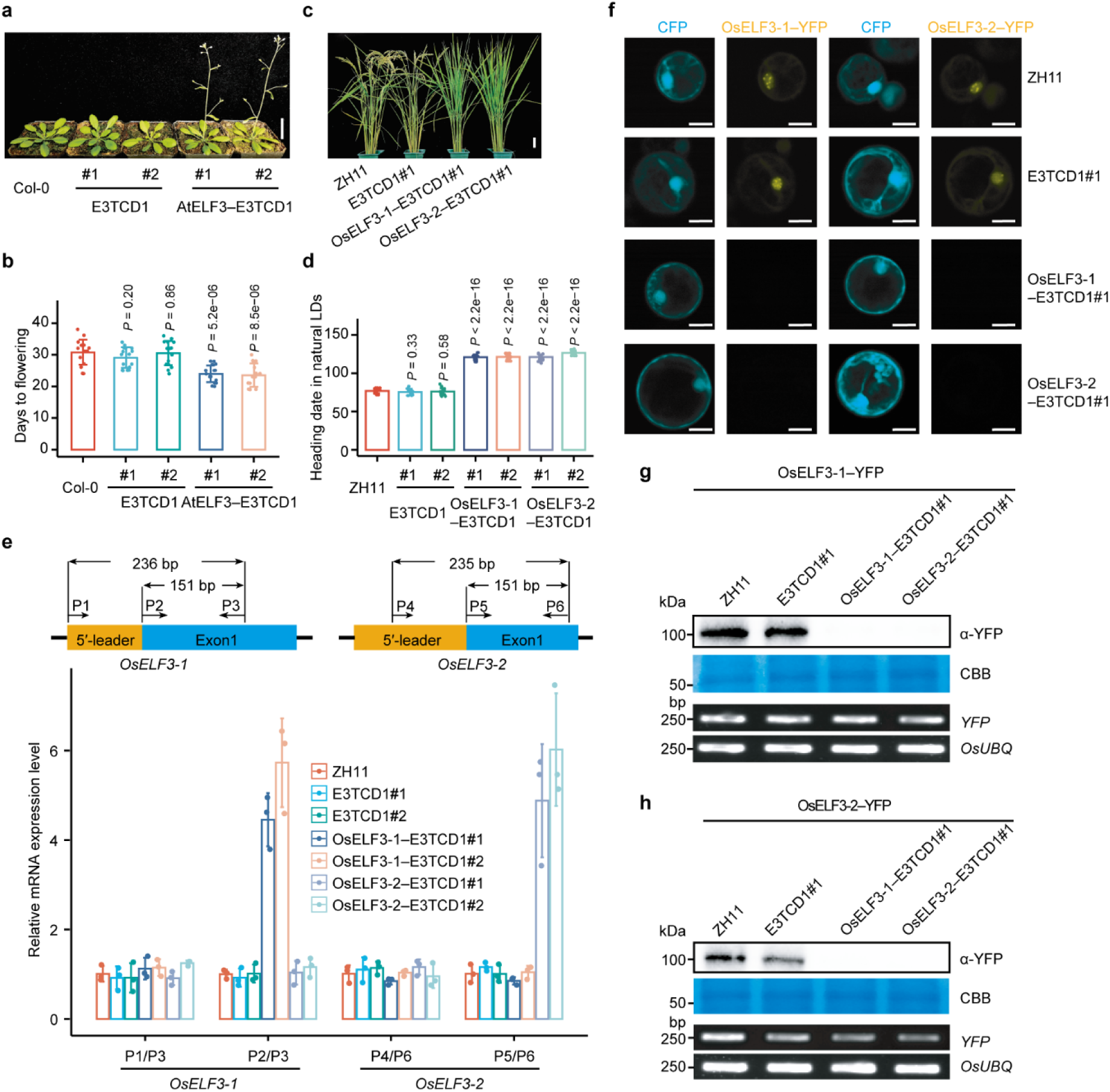
Modulating flowering times in *Arabidopsis* and rice with the TCD system. See also Extended Data Figs. 3–6. **a**, **b**, The flowering phenotype of *Arabidopsis* Col-0 plants transformed with 35S::E3TCD1 or 35S::AtELF3–E3TCD1. Two independent transgenic lines (#1 and #2) are shown. Scale bar, 3 cm. The bars (b) show the mean ± s.d. (*n* = 16) of the days to flowering after seeding, and a two-sided Student’s *t*-test was used to determine the significance. **c**, **d**, The flowering phenotype of rice Zhonghua11 (ZH11) plants transformed with 35S::E3TCD1, 35S::OsELF3-1–E3TCD1 or 35S::OsELF3-2–E3TCD1. Scale bar, 10 cm. Two independent transgenic lines (#1 and #2) were used for each construct. The bars show the mean ± s.d. (*n* = 15) of the days to heading after seeding under the natural long-day (LD) condition, and a two-sided Student’s *t*-test was used to determine the significance. **e**, Quantitative RT-PCR to show the mRNA levels of endogenous *OsELF3-1* and *OsELF3-2* in transgenic rice plants. A schematic diagram illustrating the positions of PCR primers was included. Specifically, primers P1/P3 and P4/P6 with P1 and P4 binding on the 5′-leaders were utilized to detect endogenous *OsELF3-1* and *OsELF3-2*, respectively. Additionally, PCR products generated from primers P2/P3 and P5/P6 indicated both endogenous and transgenic *OsELF3-1* and *OsELF3-2*. The bars show the mean ± s.d. after normalization to ZH11. **f**– **h**, In planta degradation assay to demonstrate the degradation of OsELF3-1 and OsELF3-2 by the TCD system by microscopic observation (f) and immunoblot analysis (g, OsELF3-1; h, OsELF3-2). The plasmids of 35S::CFP/35S::OsELF3-1–YFP or 35S::CFP/35S::OsELF3-2–YFP were co-transformed into the protoplasts prepared from the transgenic plants. Semi-quantitative RT-PCR is conducted against YFP with *NbUBQ* as the internal control (g, h). Scale bar, 10 µm.

In contrast to the inhibitory role of Arabidopsis ELF3, rice has two homologs of ELF3, with OsELF3-1 playing a more dominant role in promoting flowering than OsELF3-2 ^56–58^. We showed that both homologs exhibited high sequence similarity and interaction, as evidenced by the split-luciferase complementation assay and co-immunoprecipitation (Extended Data Fig. 6a–c). Furthermore, the co-localization assay revealed their ability to co-exist within the condensate, indicating the rice homologs are also condensation-prone proteins (Extended Data Fig. 6d, e). Given these findings, we hypothesized that OsELF3-1–E3TCD1 or OsELF3-2–E3TCD1 could target OsELF3-1 for degradation based on the design of the TCD system. Subsequently, pilot experiments conducted in *N. benthamiana* indicated that co-expression of OsELF3-1–E3TCD1 or OsELF3-2–E3TCD1 led to a reduction in YFP– OsELF3-1 (Extended Data Fig. 6f, g). Consequently, we observed a significant delay in heading date in transgenic rice lines expressing 35S::OsELF3-1–E3TCD1 or 35S::OsELF3-2–E3TCD1 compared to the control lines 35S::E3TCD1 (Fig. 3c, d). Remarkably, this observation mirrors the phenotype of plants with RNA interference of *OsELF3-1* in the Nipponbare background, as previously reported ^58^.

To validate the expression of endogenous *OsELF3-1* in transgenic plants, we conducted experiments using RT-qPCR with different primer pairs. One pair with a primer binding on the 5′-leader, which was not included in the transgene construct, while the other pair was shared by the endogenous *OsELF3-1* and the transgene (Fig. 3e). The results revealed that the transgene did not affect the levels of endogenous *OsELF3-1* mRNA, effectively ruling out the possibility of RNA interference with endogenous *OsELF3-1* in the transgenic plants (Fig. 3e). Given the unavailability of an antibody against endogenous OsELF3-1, we conducted a semi-in planta assay to demonstrate the transgene’s capability to degrade endogenous OsELF3-1 at the protein level. We introduced 35S::OsELF3-1–YFP constructs into rice protoplasts obtained from non-transformed ZH11, the control line 35S::E3TCD1, and transgenic plants expressing 35S::OsELF3-1–E3TCD1 and 35S::OsELF3-2–E3TCD1. Additionally, we introduced 35S::CFP as an internal control. Microscopic observation and immunoblot analysis targeting OsELF3-1–YFP confirmed that the TCD systems with 35S::OsELF3-1–E3TCD1 or 35S::OsELF3-2–E3TCD1 effectively degraded the target OsELF3-1 protein (Fig. 3f, g). These findings in Arabidopsis and rice underscore the TCD system in enabling controlled homologous gene functions in a species-specific manner.

### Conditional TCD in engineering crop defense

In addition to OsELF3-1, our pilot experiments revealed that OsELF3-1–E3TCD1 or OsELF3-2– E3TCD1 could facilitate the degradation of OsELF3-2. This conclusion is supported by microscopic observation and immunoblot analysis conducted in *N. benthamiana* and protoplasts prepared from 35S::OsELF3-1–E3TCD1 and 35S::OsELF3-2–E3TCD1 transgenic plants (Fig. 3f–h; Extended Data Fig. 6f, g). Intriguingly, while OsELF3-1 predominantly regulates flowering, both homologs play a vital role in suppressing resistance to *Magnaporthe oryzae*, the causal agent of rice blast disease ^59,60^. Consequently, transgenic rice lines expressing 35S::OsELF3-1–E3TCD1 and 35S::OsELF3-2– E3TCD1 exhibited reduced disease severity compared to controls (Fig. 4a, b).

**Fig. 4.**
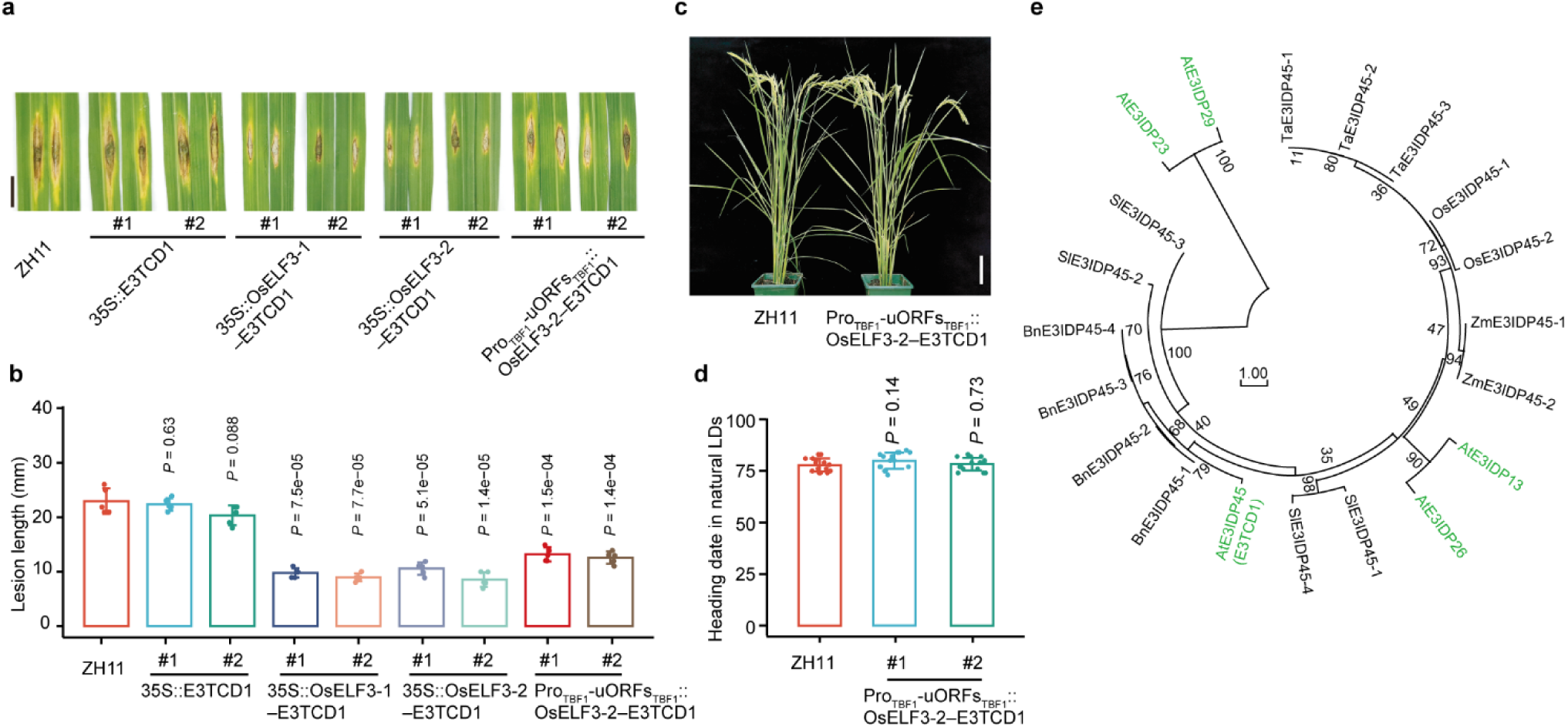
Conditional TCD with the TBF1 expression control cassette. See also Extended Data Fig. 6. **a**, **b**, Symptoms and quantification after the fungal pathogen *M. oryzae* infection in the growth chamber on rice Zhonghua11 (ZH11) plants transformed with 35S::E3TCD1, 35S::OsELF3-1–E3TCD1, 35S::OsELF3-2–E3TCD1 or Pro_TBF1_-uORFs_TBF1_::OsELF3-2–E3TCD1. Two independent transgenic lines (#1 and #2) were used for each construct. Pro_TBF1_-uORFs_TBF1_, the ‘TBF1-cassette’ consisting of the immune-inducible promoter (Pro_TBF1_) and two pathogen-responsive upstream open reading frames (uORFs_TBF1_) of the *Arabidopsis TBF1* gene. Scale bar, 1 cm. The bars show the mean ± s.d. (*n* = 5) of the lesion length, and a two-sided Student’s *t*-test was used to determine the significance. **c**, **d**, The flowering phenotype of rice Zhonghua11 (ZH11) plants transformed with Pro_TBF1_-uORFs_TBF1_::OsELF3-2–E3TCD1. Scale bar, 10 cm. Two independent transgenic lines (#1 and #2) were used for each construct. The bars show the mean ± s.d. (*n* = 15) of the days to heading after seeding under the natural long-day (LD) condition, and a two-sided Student’s *t*-test was used to determine the significance. **e**, The phylogenetic tree to show the homologs of E3TCD1 in *Arabidopsis thaliana* (At), *Oryza sativa* (Os), *Zea mays* (Zm), *Triticum aestivum* (Ta), *Brassica napus* (Bn), and *Solanum lycopersicum* (Sl).

To mitigate the pleiotropic effects associated with continuous degradation of OsELF3-1 and OsELF3-2, such as late flowering, we employed the TBF1 cassette to control the TCD system precisely. This cassette allows transcriptional and translational control of target genes by incorporating an immune-inducible promoter (Pro_TBF1_) and two pathogen-responsive upstream open reading frames (uORFs_TBF1_) of the *TBF1* gene ^61,62^. Remarkably, transgenic plants with Pro_TBF1_-uORFs_TBF1_::OsELF3-2–E3TCD1 showed no alteration in flowering time compared to non-transformed ZH11 plants, indicating stringent control of *OsELF3-2* expression by the TBF1 expression control cassette (Fig. 4c, d). However, these plants exhibited enhanced resistance to *M. oryzae* (Fig. 4a, b). These findings underscore the potential of the TCD system coupled with expression-control elements for conditionally targeted protein degradation, offering practical utility in engineering desirable agronomic traits for genes with pleiotropic effects.

## Discussion

Modern agriculture faces many challenges, such as a rapidly growing global population, decreasing arable land, unpredictable climate patterns, and environmental pollution. To foster sustainable agriculture, there’s an urgent need for the rapid development of high-yield food and fiber crops resilient to biotic and abiotic stresses. Genetic engineering emerges as a potent tool for swiftly breeding superior cultivars, adept at navigating these challenges by modulating the expression of essential agronomic trait genes across various levels. In this context, we introduce the TCD system, enabling the degradation of endogenous proteins via transgene expression, complementing genetic techniques that manipulate DNA and RNA in plants. These findings will deepen our understanding of protein functionalities in fundamental research and hint at practical applications, potentially as a pivotal gene therapeutic strategy for optimizing crop performance and combating human diseases.

The success of the TCD system largely depends on identifying an appropriate E3 ligase, referred to as E3TCDs, which can induce self-degradation either independently or when fused with a target protein (X protein). Our study demonstrates the functional efficacy of Arabidopsis E3TCD1 in targeting specific TFs for degradation in various expression systems, including Arabidopsis, *N. benthamiana*, and rice. Specifically, E3TCD1 successfully degraded 10 out of 12 tested TFs or transcriptional regulators. However, it failed to degrade some TFs despite their similar subcellular localization patterns to the degradable TFs (Extended Data Fig. 2a). This suggests that the X–E3TCD1 in the TCD system might interfere with the recognition of some endogenous TF or the recruitment of other UPS components, resulting in varying degradation efficiencies. Adjusting the conformation between the X protein and E3TCD1 might be beneficial to address this issue. Additionally, reducing the length of E3TCD1 from its current 666 amino acids may minimize conformational or structural interference. A more promising approach would be to identify additional E3TCDs to broaden the recognition spectrum of the X protein. Although our current study screened candidate E3TCDs from a subset of E3 ligases predicted to be IDPs, we discovered that the primary characteristic of effective E3TCDs is their propensity for self-degradation. We identified numerous E3TCD1 homologs across various plant species (Fig. 4e). These data suggest the potential for discovering more E3TCDs to expand the substrate recognition spectrum, enhancing the versatility and efficacy of the TCD system. Therefore, a large-scale screening of E3 ligases from various species might be necessary.

The next question concerns the situations where the TCD system could be considered for genetic manipulations. We believe that for reverse genetics with known genes, the TCD system could be applied in scenarios where CRISPR interference (CRISPRi) or RNA interference (RNAi) methods are currently used once the positive results from our pilot experiments. Based on our study of OsELF3-1/2 homologs in rice, the TCD system demonstrates superiority in studying homologous genes with higher identity at the protein level than at the nucleotide level. Unlike CRISPRi or RNAi, which require the expression of multiple gRNAs or RNA fragments to reduce the expression of homologous genes, the TCD system can target and degrade multiple homologs simultaneously using the TCD with a single gene. For example, in our study, the TCD system effectively reduced the expression of both OsELF3-1 and OsELF3-2 using the TCD with either OsELF3-1 or OsELF3-2, highlighting its efficiency and potential advantages in genetic manipulations involving homologous genes. This advantage may become more pronounced in forward genetic screening, particularly given the widespread occurrence of gene redundancy in plant genomes ^63^.

In crop engineering, it’s crucial to focus on how to express the transgene within the TCD system. This is important because most genes have multiple functions, leading to potential pleiotropic effects. Therefore, TCD must be precisely regulated to control when, where, and how strongly the gene is expressed. Our study used the TBF1 cassette to control TCD timing, successfully avoiding pleiotropic effects on flowering time caused by OsELF3-1. In addition, adjusting flowering times in rice cultivars could enable rapid adaptation to different light conditions across latitudes. TCD-mediated degradation of OsELF3-1 significantly changed flowering times (Fig. 3c, d). This highlights the possibility of selecting expression intensity to fine-tune flowering time to meet the different requirements of the breeding programs on flowering time. Thus, future research could develop quantitative TCD methods to achieve gradient degradation of specific proteins, resulting in variable phenotypes. This variability could benefit breeding elite cultivars tailored to specific environments, considering factors like light intensity, water availability, temperature fluctuations, and soil composition.

Our study has a limitation in explaining the scale at which TCD works on cleaning condensates away. This is because liquid-liquid phase separation facilitates frequent exchange between the soluble fraction and the condensate. Therefore, it is possible that TCD may lead to direct degradation of either the condensate or the smaller-sized complex in the soluble fraction that is continuously exchanged from the condensate. In either scenario, there will be a notable decrease in the amount of the targeted proteins and the corresponding condensates, as observed. Certain condensates may turn into harmful protein condensates or solidified fiber structures. These harmful condensates have been recently found to compromise plant disease tolerance, and they are associated with neurodegenerative diseases in humans, such as amyotrophic lateral sclerosis (ALS) and Alzheimer’s disease ^30,64^. In such cases, the TCD system may effectively eliminate these harmful condensates. Notably, during the peer review of this work, a similar strategy called ‘RING-Bait’ was reported. This approach combines an aggregating protein sequence with the E3 ubiquitin ligase TRIM21 to specifically degrade tau aggregates, ultimately reducing tau pathology and improving motor function in mice ^65^. Additionally, another strategy, known as the ‘RING-nanobody degrader,’ has been developed. Unlike RING-Bait and our TCD system, this approach fuses the RING domain of TRIM21 to a tau-targeting nanobody, also demonstrating therapeutic potential for tau-associated disorders ^66^. Our TCD system targets distinct properties of endogenous proteins, specifically TFs, which are widely linked to various human diseases. As a result, our TCD system holds therapeutic potential for a range of diseases driven by endogenous TFs. Furthermore, our approach begins with de novo mining of unknown E3 ligases, which could help identify E3 candidates suitable for gene therapy in human diseases caused by other types of endogenous proteins. With the rapid advancements in targeted protein degradation strategies ^67–69^, it is exciting to envision their application in plants, where they could significantly accelerate basic research and enhance crop performance, paving the way for an exciting future.

## Methods

### Plant growth and transformation

All rice lines in this research belong to the *O. sativa* subsp. *japonica* variety Zhonghua 11 (ZH11). Rice plants for phenotypic assessments were cultivated in the experimental field at Huashan in Wuhan, China (30.3°N, 114.3°E) under natural long-day conditions (LDs). Rice plants for *Magnaporthe oryzae* infection were grown in a greenhouse at 26°C under a photoperiod of 12 hours of light and 12 hours of darkness with 90% relative humidity. All Arabidopsis lines in this study originate from the Columbia-0 (Col-0) background. Arabidopsis and *N. benthamiana* plants were grown in a greenhouse at 23°C under a photoperiod of 16 hours of light and 8 hours of darkness with 55% relative humidity. The genetic transformation was performed in the background of ZH11 and Col-0 as described previously ^22,70^.

### Plasmid construction

The full-length coding sequences of *AtELF3* (At2G25930.1) and *E3IDPs* (Supplementary Table 1) were amplified from Col-0 cDNA. The CDSs of *OsTB1* (LOC_Os03g49880.1), *OsELF3-1* (LOC_Os06g05060.1), *OsELF3-2* (LOC_Os01g38530.1), and *OsPLPs* (Supplementary Table 2) were amplified from ZH11 cDNA. All sequences were fused with the indicated N or C terminal tags. All constructs were confirmed through conoly PCR and DNA sequencing. All primers and plasmids are listed in Supplementary Table 4.

### Immunoblot analysis

In immunoblot experiments, protoplasts, *N. benthamiana*, and rice tissues were collected and ground in liquid nitrogen using a steel bead. Subsequently, 200 μL of lysis buffer (50 mM Tris, pH 7.5, 150 mM NaCl, 0.1% Triton X-100, 0.2% Nonidet P-40, and a protease inhibitor cocktail (Thermo Fisher, Cat#A32955; one tablet per 10 mL) was added to each 100 mg of ground tissue samples. The lysates were incubated at 4°C for 10 minutes, followed by centrifugation at 12,000 × g to collect the supernatant. Proteins in the supernatant were denatured by the addition of 1× SDS-loading buffer and heating at 100°C for 10 minutes. The samples were then subjected to Western blotting using anti-GFP (mouse mAb, Abclonal) or anti-HA (mouse mAb, Abclonal) antibodies.

### Transient expression in *N. benthamiana*

We employed *Agrobacterium* strain GV3101 (pMP90) for binary vector delivery. For transient expression in *N. benthamiana*, *Agrobacterium* GV3101 carrying each construct was cultured in LB medium supplemented with kanamycin (50 μg ml^−1^), gentamycin (50 μg ml^−1^) and rifampicin (25 μg ml^−1^) at 28℃ overnight. The agrobacterial cells were resuspended in the infiltration buffer (10 mM 2-(N-morpholino) ethanesulfonic acid, 10 mM MgCl_2_, 200 μM acetosyringone) at OD_600 nm_ = 0.1 after centrifugation at 2,600 × g for 5 min and incubated at room temperature for 1 h.

For microscopic observation in the pilot experiments, *Agrobacterium* containing 35S::CFP and *Agrobacterium* containing 35S::X–YFP or 35S::YFP–X were mixed with *Agrobacterium* containing 35S::E3TCD1 or 35S::X–E3TCD1 at a 1:1:1 ratio. The mixed suspension was infiltrated into three-week-old *N. benthamiana* plants. After 48 hours of expression, leaf discs were excised for microscopic observation. CFP and YFP fluorescence were visualized using Leica TCS SP8 upright microscopy, with excitation provided by 448 nm and 514 nm lasers, respectively.

For split luciferase complementation assays, *Agrobacterium* containing the indicated pairwise vectors were mixed at a 1:1 ratio. Negative controls, 35S::HA–nLUC or 35S::nYFP–cLUC, were included. Both the test and control groups were infiltrated into the same leaf of *N. benthamiana* and allowed to express for 48 hours. Subsequently, 1 mM luciferin was sprayed onto the leaves, and imaging was conducted using a CCD-camera-equipped box at an exposure time of 20 min under darkness.

For the in vivo analysis of protein degradation in *N. benthamiana*, leaves were infiltrated with Agrobacterium tumefaciens harboring the construct 35S::PLP308–YFP. Following a 24-hour incubation period, the same leaf areas were subsequently infiltrated with Agrobacterium containing the est::PLP308–E3TCD1 construct. After an additional 24 hours of incubation, the leaves were sectioned into three groups and treated with 10 mM MgCl_2_, 50 μM β-estradiol, or a combination of 50 μM β-estradiol and 50 μM MG132. Samples were collected at 0, 6, 12, and 24 hours post-treatment for immunoblot analysis to quantify PLP308–YFP protein levels. In a parallel experiment, leaves were co-infiltrated with Agrobacterium containing 35S::PLP308–YFP and 35S::CFP. These leaves were then infiltrated with Agrobacterium harboring the est::PLP308–E3TCD1 construct, followed by treatment with 50 μM β-estradiol as described. Leaf samples were collected at 0 and 12 hours post-treatment for fluorescence microscopy to observe protein localization and degradation.

For co-immunoprecipitation (co-IP) analysis in *N. benthamiana*, Agrobacterium containing 35S::YFP–OsELF3-1 was mixed in equal amounts with Agrobacterium containing 35S::HA–OsELF3-2 or 35S::HA–nLUC. The mixed suspension was infiltrated into three-week-old N. benthamiana plants. After 48 hours of expression, 1 g of leaf tissue was collected from each sample for protein extraction, following the method described previously. From the 2 mL of total extracted protein, 90 μL was taken as input, and the remaining protein was incubated with 100 μL GFP-Trap_A beads on a rotating shaker for 1 hour at 4°C. The beads were then collected using a magnetic separator, the supernatant was discarded, and the beads were washed with 1 mL of washing buffer (50 mM Tris, pH 7.5, 150 mM NaCl, 1 mM PMSF) for 1 minute. This washing step was repeated three times, after which the supernatant was discarded. Finally, the bead-bound proteins were mixed with 100 μL of 1× SDS-loading buffer, and the 90 μL input samples were mixed with 10 μL of 10× SDS-loading buffer. Both sets of samples were then heated at 100°C for 10 minutes. Subsequently, the samples were subjected to immunoblot analysis.

### Pathogen infection

Isolates of *Magnaporthe oryzae* strain RB22 were incubated on an oatmeal medium (3% [w/v] oats and 1.5% [w/v] agar) The cultures were incubated at 26°C under low light conditions for 18 days. Following incubation, spores were harvested by washing the plates with a small volume of 0.1% (v/v) Tween 20. The spore suspensions were subsequently filtered through Miracloth and diluted with 0.1% (v/v) Tween 20 to achieve a final concentration of 5.0 × 10⁵ spores ml⁻¹. For the infection assay, wounds were created on the abaxial surface of rice leaves. A 10 µL aliquot of the spore suspension was subsequently applied to each wound site. The inoculated leaves were then covered with cellophane tape to maintain humidity. After a ten-day incubation period, disease severity was assessed by measuring the length of the lesions on the rice leaves.

### In vitro phase separation assay

The assay was conducted following the method described previously ^21^. The MBP–YFP–E3IDP45 fusion protein was expressed in *E. coli* (DE3) cells. Post cell lysis and purification, the MBP tag was enzymatically cleaved using 10 μg ml^-1^ Xa factor protease (NEB, Cat#P8010) at 23°C overnight. To induce phase separation, PEG8000 (Sigma, Cat#25322-68-3) was added to the indicated final concentrations. Both MBP–YFP–E3IDP45 and YFP–E3IDP45 were then coated onto confocal flat dishes (Solarbio) and observed using a Leica TCS SP8 upright microscope equipped with a 10× objective lens and a 514 nm laser for excitation. FRAP was performed by bleaching the region of interest with a 514 nm laser at 80% power intensity, applied twice. The recovery of YFP fluorescence was monitored at intervals of 1.3 seconds post-bleaching.

### Transient expression in the rice protoplast

To transiently express genes in rice protoplasts, dehulled rice seeds were disinfected using a 2% hypochlorous acid solution. The disinfected seeds were then planted on 1/2 MS medium and incubated in a light chamber at 28°C under a photoperiod of 14 hours of light and 10 hours of darkness with 60% relative humidity. After 14 days, approximately 40 uniformly grown seedlings were selected, and seedling stem were excised into small segments approximately 1 mm in length. These segments were incubated in 20 ml of enzyme solution (1.5% Cellulose RS (Yacult Pharmaceutical), 0.75% Macerozyme R-10 (Yacult Pharmaceutical), 0.6M Mannitol, 1 mM CaCl_2_, 0.1% BSA, 10 mM MES (pH 5.7)) on a rotary shaker at 40 rpm and 25°C for 5 hours to facilitate enzymatic digestion. The resulting digestion mixture was filtered through a 40 μm nylon mesh to eliminate plant debris. Protoplasts were harvested by centrifugation at 100 × g, and the supernatant was discarded. The protoplasts were subsequently washed with W5 solution (154 mM NaCl, 5 mM KCl, 125 mM CaCl_2_, and 2 mM MES, pH 5.7) and resuspended in MMG solution (0.6 M mannitol, 15 mM MgCl_2_, and 4 mM MES, pH 5.7) at a concentration of 1 × 10^7^ cells ml^-1^. For transformation, 5 μg of the desired plasmid was added to 100 μl of the protoplast suspension, followed by incubation with an equal volume of PEG-CaCl_2_ solution (0.6 M mannitol, 100 mM CaCl_2_, and 40% PEG4000) for 15 minutes. The reaction was terminated by the addition of 1 ml W5 solution, and the protoplasts were collected by centrifugation at 100 × g. The transformed protoplasts were then transferred to 1 ml WI solution (0.6 M mannitol, 4 mM KCl, and 4 mM MES, pH 5.7) and incubated in the dark at 28°C for 16 hours. Following the incubation period, the protoplasts were subjected to microscopic observation and protein extraction.

### Semi-in vivo protein degradation assay

In the semi-in vivo protein degradation assay, 5 μg of the 35S::CFP plasmid was combined with 5 μg of either the 35S::YFP–OsELF3-1 or 35S::YFP–OsELF3-2 plasmid. These plasmid mixtures were subsequently introduced into 100 μl of rice protoplasts, utilizing the transformation method previously described. The protoplasts were isolated from non-transformed ZH11, control line 35S::E3TCD1, and transgenic lines expressing 35S::OsELF3-1–E3TCD1#1 or 35S::OsELF3-2–E3TCD1#1. Following a 16-hour transformation period, the protoplasts were subjected to confocal microscopy and immunoblot analysis.

### RT-qPCR and RT-PCR

Total RNA was isolated from 100 mg of leaf tissue using 1 mL of TRIzol (Vazyme, Cat#R401-01) following the manufacturer’s instructions. Subsequently, reverse transcription was carried out using the HiScript III 1st Strand cDNA Synthesis Kit (Vazyme, Cat#R312-01). Real-time quantitative PCR (qPCR) assays were performed using Taq Pro Universal SYBR qPCR Master Mix (Vazyme, Cat#Q712-02). *OsActin1* was utilized as internal control for rice. RT-PCR was conducted using Phanta Max Super-Fidelity DNA Polymerase (Vazyme, Cat#P505-d1). In experiments involving rice protoplasts and Nicotiana benthamiana, *OsUbiquitin* (*OsUBQ*, LOC_Os03g13170) and *NbUbiquitin* (*NbUBQ*, AY912494) served as internal controls, respectively. Detailed information on primers and plasmids can be found in Supplementary Table 4.

### Phylogenetic tree analysis

Phylogenetic analysis was conducted using a diverse range of species including *Arabidopsis thaliana*, *Oryza sativa*, *Zea mays*, *Triticum aestivum*, *Brassica napus*, and *Solanum lycopersicum*. Predicted ELF3 protein sequences were retrieved from the Plant orthologs of AtELF3 in TAIR (https://www.arabidopsis.org/), while predicted E3IDP45 protein sequences were obtained from the Protein BLAST tool available on the National Center for Biotechnology Information (NCBI) database (https://www.ncbi.nlm.nih.gov/). The BLAST search was performed using default parameters. Detailed information on predicted protein can be found in Supplementary Table 4. For construction of the phylogenetic tree, each protein sequence was aligned using ClustalW. Subsequently, 1,000 bootstrap replicates were generated for evolutionary analysis using the Neighbor-Joining method. These analyses were carried out using MAGA11^71^.

## Data availability

All materials are available from the corresponding author upon request. All data generated or analysed during this study are included in this article and its Extended Data.

## Statistical methods

Our comprehensive analysis employed rigorous statistical methodologies using GraphPad Prism 8 or RStudio. We assumed a normal distribution for parametric statistics when the Shapiro-Wilk test yielded a *P*-value greater than 0.05. For comparisons between two groups, we used a two-sided Student’s *t*-test. E3 screening and in vitro phase separation experiments have only been performed once. Unless stated otherwise, ’*n*’ represents biological replicates, and all experiments were conducted independently at least three times. Additional statistical parameters are outlined in the Figure legends for further elucidation.

## Author Contributions

G.X. and M.L. designed the experiment. M.L., S.Z. R.N. and G.X. analyzed the data and prepared the figures. S.Z. and R.N. performed all bioinformatics analysis. M.L. performed the experiments with the help of H.D., Q.W, J.L. and Z.Wang. Y.T., Y.N. and M.Y. helped with data analysis. G.X. and M.L. wrote the manuscript with input from all authors.

## Acknowledgements

This study was supported by grants from the National Key R&D Program of China (2023ZD04073) to G.X., the Major Project of Hubei Hongshan Laboratory (2022hszd016) and the National Natural Science Foundation of China and the Key Research and Development Program of Hubei Province (2022BFE003) to G.X.

## Supplemental Information

Supplemental Information includes 6 figures and 4 tables, and can be found with this article online at XXX.

## Supplementary Information

**Extended Data Fig. 1.**
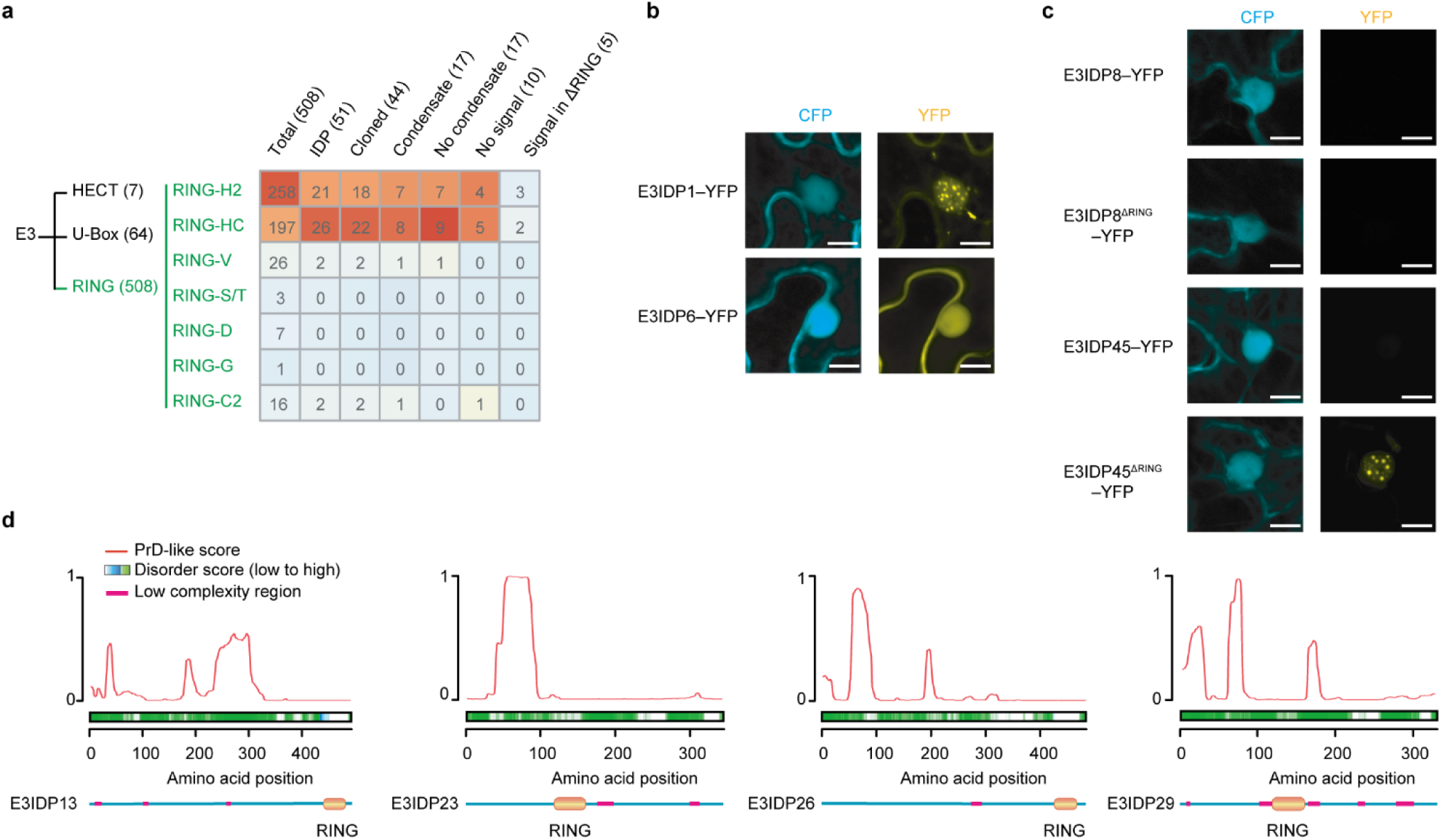
Screening E3 ligases for the TCD system. Related to Fig. 1. **a**, The heatmap to show the number of E3 ligases in each subfamily for the screening. A total of 508 E3 ligases from the reference dataset were analyzed using PLAAC to predict their status as intrinsically disordered proteins (IDPs). Of these, 51 were identified as potential IDPs (E3IDPs). Among these E3IDPs, 44 were successfully cloned and expressed in *N. benthamiana* as YFP fusion proteins (E3IDP–YFP). Microscopic analysis of these constructs revealed that 17 E3IDPs formed visible condensates, 17 remained soluble, and 10 exhibited no detectable fluorescence signals. Notably, for 5 of the E3IDPs that initially showed no signal, deletion of the RING domain (ΔRING) restored YFP fluorescence within condensates. **b**, Exemplifying E3IDPs showing visible condensates (E3IDP1) or existing in soluble fractions (E3IDP6). Scale bar, 10 µm. **c**, Exemplifying E3IDPs showing no detectable signals. Out of the E3IDPs analyzed, 5 of them still displayed no detectable signals even after the predicted RING domain was removed (e.g., E3IDP8). However, in the case of the other 5 E3IDPs, the signals were restored upon removing the predicted RING domain, which was considered the E3IDP candidates for the TCD (e.g., E3IDP45). Scale bar, 10 µm. For microscopic observation, 35S::CFP serves as the control (b, c). **d**, Prediction of the five TCD candidate E3IDPs as an intrinsically disordered protein (IDP) using PLAAC and D2P2 algorithms indicated by Prion domain (PrD)-like score and disorder score, respectively. The bottom section displays low complexity regions and the RING domain as annotated by the website SMART. E3IDP45 is shown in Fig. 1b.

**Extended Data Fig. 2.**
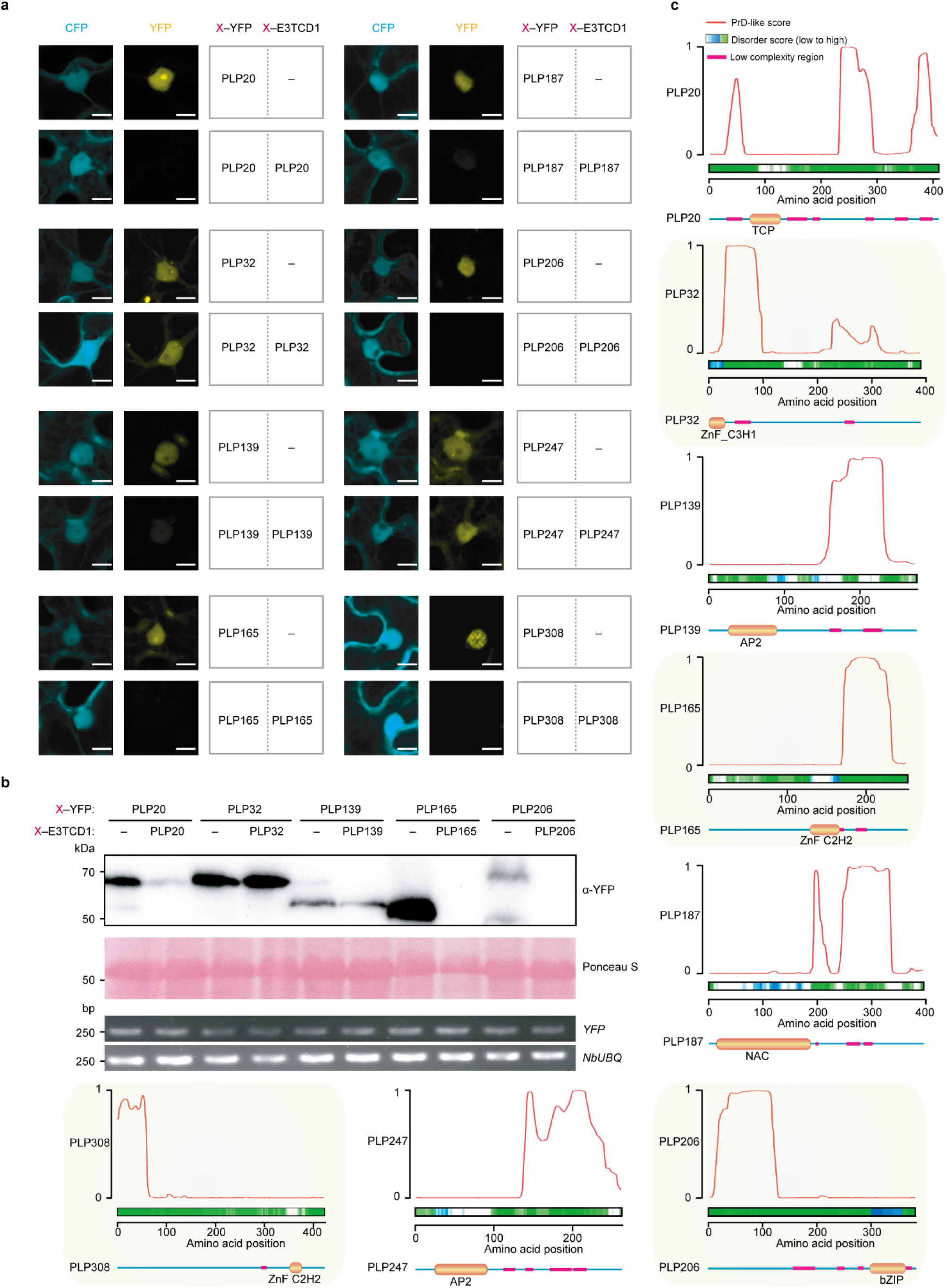
Validating the TCD system using transcription factors. Related to Fig. 2. **a**, **b**, Pilot experiments to show the degradation results of eight transcription factors by the respective TCD system through microscopic observation (a) and immunoblot analysis (b). Each X–YFP was co-expressed with X–E3TCD1 or E3TCD1 alone in *N. benthamiana*. Six X–YFPs were successfully detected in the immunoblot analysis with PLP308 shown latter. For microscopic observation, 35S::CFP serves as the control (a). Semi-quantitative RT-PCR was conducted against YFP with *NbUBQ* as the internal control (b). Scale bar, 10 µm. **c**, Prediction of the eight transcription factors as an intrinsically disordered protein (IDP) using PLAAC and D2P2 algorithms indicated by Prion domain (PrD)-like score and disorder score, respectively. The bottom section displays low complexity regions and the TF domain as annotated by the website SMART.

**Extended Data Fig. 3.**
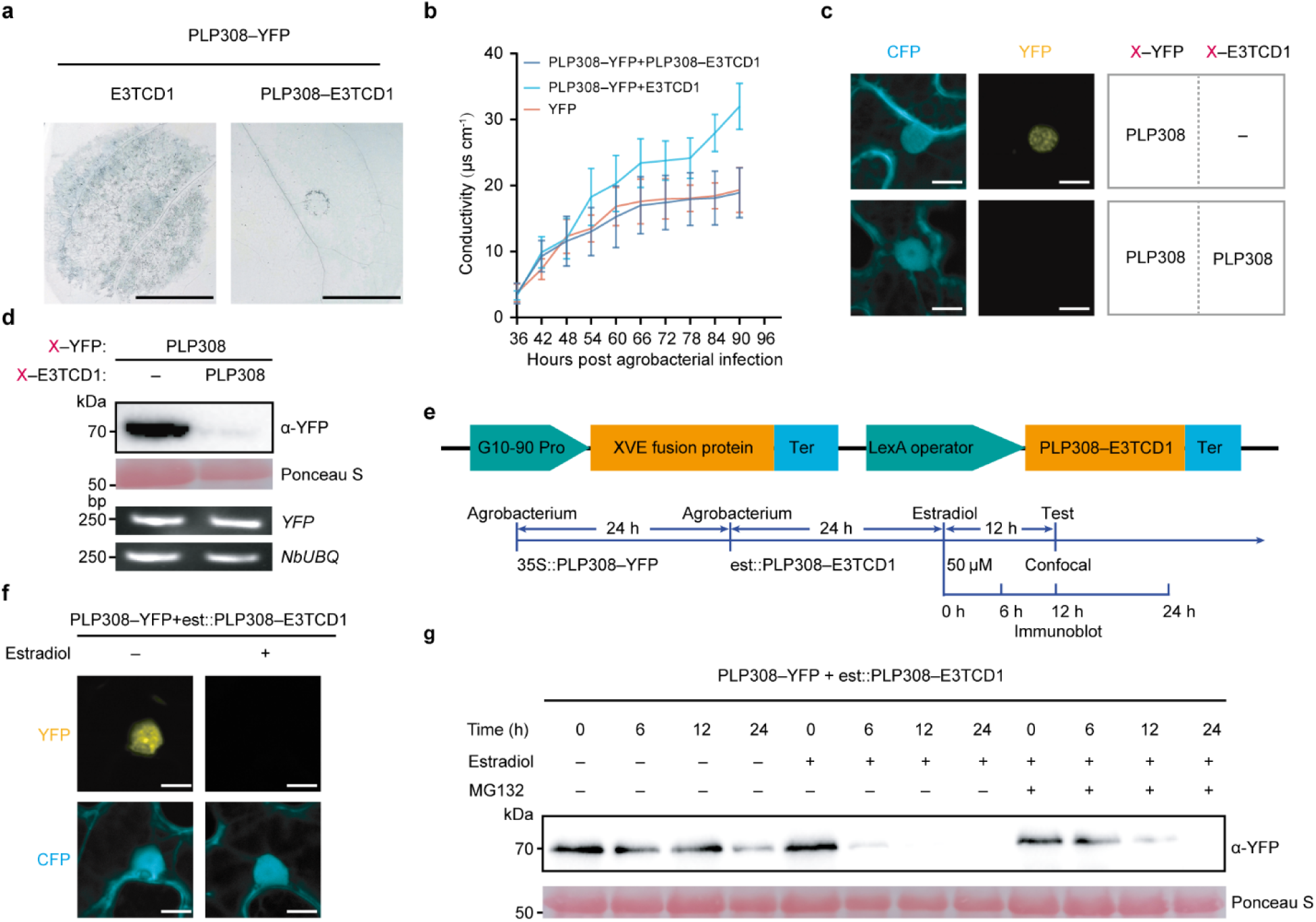
The TCD system uncovers the role of PLP308 in promoting cell death. Related to Fig. 3. **a**, Trypan blue staining to show PLP308–YFP-triggered cell death without (E3TCD1) or with the TCD (PLP308–E3TCD1). Macroscopic cell death appears after transient expression of PLP308–YFP in *N. benthamiana* for 72 hours. **b**, Ion leakage to show PLP308–YFP-triggered cell death. YFP alone was used as a control. Leaf discs were used for measuring ion leakage after transient expression of PLP308–YFP in *N. benthamiana* for 36 hours. The points and error bars show the mean ± s.d. of ion concentration (*n* = 4; 6 discs each). **c**, **d**, Pilot experiments to show the degradation of PLP308–YFP by PLP308–E3TCD1 through microscopic observation (c) and immunoblot analysis (d) after transient expression in *N. benthamiana* for 48 hours before the appearance of macroscopic cell death. **e**, The schematic of the β-estradiol-inducible system. In this setup, PLP308–YFP expression was under the control of the CaMV 35S constitutive promoter (35S::PLP308–YFP), while PLP308–E3TCD1 expression was regulated by β-estradiol (est::PLP308–E3TCD1). Following the infiltration of agrobacterium containing 35S::PLP308–YFP for 24 hours, agrobacterium containing est::PLP308– E3TCD1 was infiltrated for another 24 hours. The treatment with β-estradiol triggers the movement of the XVE fusion protein into the nucleus. Once inside the nucleus, XVE binds to the LexA operon, initiating the transcription of PLP308– E3TCD1.Tests by confocal observations and immunoblot analysis were performed at the indicated time points after est trea. G10-90 Pro, the strong constitutive promoter to drive the transcription of XVE fusion protein. XVE, the fusion of the DNA-binding domain of the bacterial repressor LexA (X), the acidic transactivating domain of VP16 (V) and the regulatory region of the human estrogen receptor (E; ER). Ter, transcription terminators. **f**, Degradation of PLP308–YFP by PLP308– E3TCD1 controlled by the β-estradiol-inducible system through microscopic observation. Agrobacterium containing 35S::CFP was co-infiltrated with agrobacterium containing 35S::PLP308–YFP, serving as the control. Scale bar, 10 µm. **g**, Degradation of PLP308–YFP by PLP308–E3TCD1 controlled by the β-estradiol-inducible system through immunoblot analysis. In vitro protein degradation assay was performed by collecting samples after β-estradiol treatment for 0, 6, 12, and 24 hours in the absence or presence of MG132, the inhibitor of the 26S proteasome.

**Extended Data Fig. 4.**
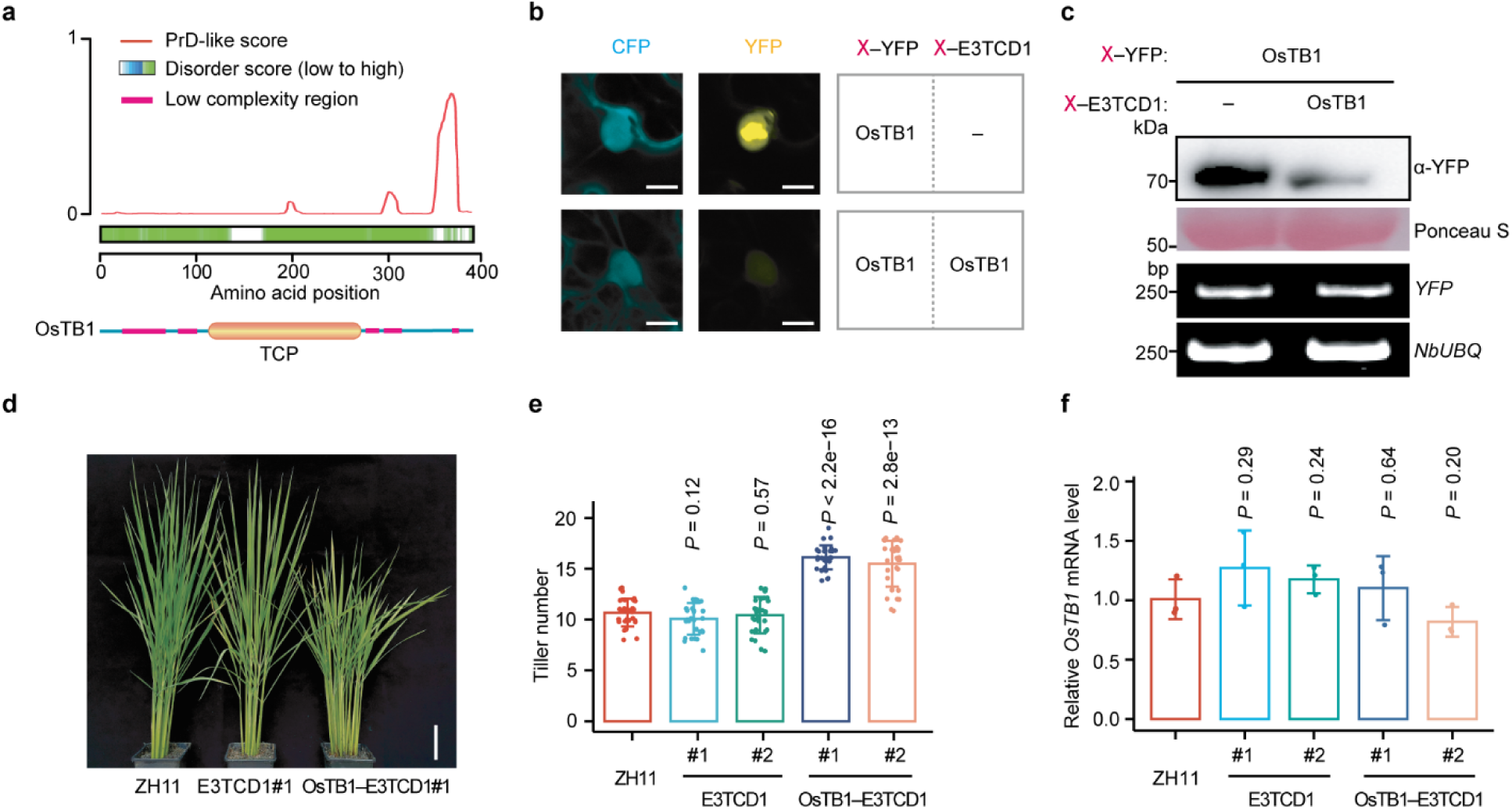
Modulating rice tiller numbers by the TCD system in rice. Related to Fig. 3. **a**, Prediction of OsTB1 as an intrinsically disordered protein (IDP) using PLAAC and D2P2 algorithms indicated by Prion domain (PrD)-like score and disorder score, respectively. The bottom section displays low complexity regions and the RING domain as annotated by the website SMART. **b**, **c**, Pilot experiments to show the degradation of OsTB1 by the TCD system through microscopic observation (b) and immunoblot analysis (c). For microscopic observation, 35S::CFP serves as the control (b). Semi-quantitative RT-PCR is conducted against YFP with *NbUBQ* as the internal control (c). Scale bar, 10 µm. **d**, **e**, The rice tiller numbers in rice Zhonghua11 (ZH11) plants transformed with 35S::E3TCD1 or 35S::OsTB1–E3TCD1. Two independent transgenic lines (#1 and #2) were used for each construct. The bars show the mean ± s.d. (*n* = 30) of tiller numbers, and a two-sided Student’s *t*-test was used to determine the significance. Scale bar, 10 cm. **f**, Quantitative RT-PCR to show the relative endogenous *OsTB1* mRNA levels in the transgenic lines. Primers binding to the untranslated region, which was not included in the OsTB1–E3TCD1 transgene, were used to amplify the endogenous *OsTB1* gene. The bars show the mean ± s.d. (*n* = 3) after normalization to ZH11, a two-sided Student’s *t*-test was used to determine the significance.

**Extended Data Fig. 5.**
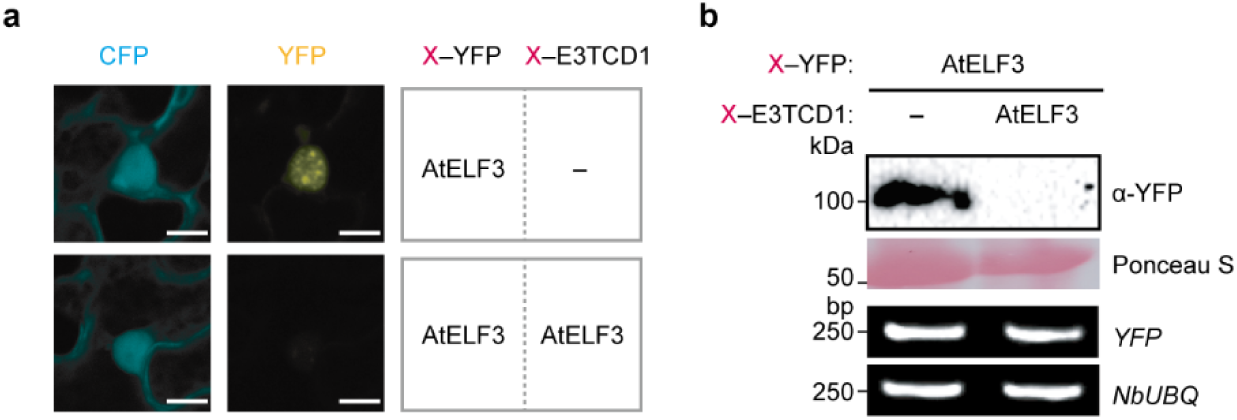
Pilot experiments to show the degradation of AtELF3 by the TCD system. Related to Fig. 3. **a**, **b**, Through microscopic observation (a) and immunoblot analysis (b). For microscopic observation, 35S::CFP serves as the control (a). Semi-quantitative PCR is conducted against YFP with *NbUBQ* as the internal control (b). Scale bar, 10 µm.

**Extended Data Fig. 6.**
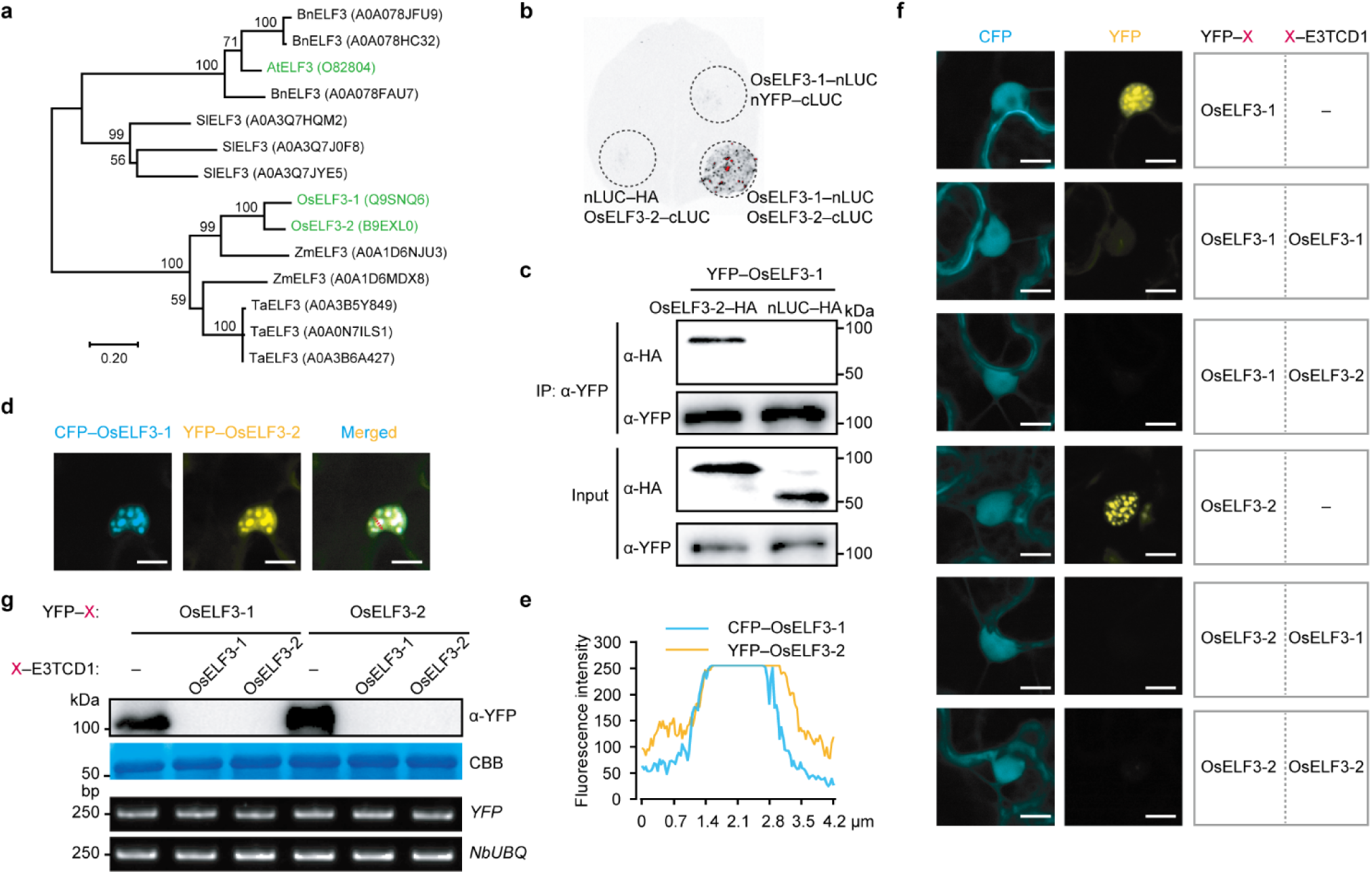
Degradation of OsELF3-1 and OsELF3-2 by the TCD system. Related to Figs. 3 and 4. **a**, The phylogenetic tree to show the number and their relationship among ELF3 homologs in *Arabidopsis thaliana* (At), *Oryza sativa* (Os), *Zea mays* (Zm), *Triticum aestivum* (Ta), *Brassica napus* (Bn), and *Solanum lycopersicum* (Sl). **b**, Split luciferase complementation assay (SLCA) to show the interaction of OsELF3-1 and OsELF3-2 in *N. benthamiana*. nLUC and cLUC, the N and C terminal domains of the firefly luciferase, respectively. nLUC–HA and nYFP–cLUC were used as a negative control. **c**, Co-immunoprecipitation to show the interaction of OsELF3-1 and OsELF3-2. YFP–OsELF3-1 was co-expressed alongside either OsELF3-2–HA or nLUC–HA in *N. benthamiana* for a duration of 48 hours. The resulting OsELF3-1 complex was then subjected to immunoprecipitation using GFP-trap beads. Following this, immunoblot analysis was conducted utilizing antibodies specific to YFP or HA. **d**, **e**, The co-localization assay between CFP–OsELF3-1 and YFP–OsELF3-2 in *N. benthamiana*. The fluorescence intensity in (e) was measured along the red arrow in (d). Scale bars, 10 µm. **f**, **g**, Pilot experiments to show the degradation of OsELF3-1 and OsELF3-2 by the TCD system through microscopic observation (f) and immunoblot analysis (g). For microscopic observation, 35S::CFP serves as the control (f). Semi-quantitative RT-PCR is conducted against YFP with *NbUBQ* as the internal control (g). Scale bar, 10 µm. CBB, Coomassie brilliant blue.

